# Rapid invisible frequency tagging reveals nonlinear integration of auditory and visual information

**DOI:** 10.1101/2020.04.29.067454

**Authors:** Linda Drijvers, Ole Jensen, Eelke Spaak

## Abstract

During communication in real-life settings, the brain integrates information from auditory and visual modalities to form a unified percept of our environment. In the current magnetoencephalography (MEG) study, we used rapid invisible frequency tagging (RIFT) to generate steady-state evoked fields and investigated the integration of audiovisual information in a semantic context. We presented participants with videos of an actress uttering action verbs (auditory; tagged at 61 Hz) accompanied by a gesture (visual; tagged at 68 Hz, using a projector with a 1440 Hz refresh rate). Integration difficulty was manipulated by lower-order auditory factors (clear/degraded speech) and higher-order visual factors (congruent/incongruent gesture). We identified MEG spectral peaks at the individual (61/68 Hz) tagging frequencies. We furthermore observed a peak at the intermodulation frequency of the auditory and visually tagged signals (f_visual_ - f_auditory_ = 7 Hz), specifically when lower-order integration was easiest because signal quality was optimal. This intermodulation peak is a signature of nonlinear audiovisual integration, and was strongest in left inferior frontal gyrus and left temporal regions; areas known to be involved in speech-gesture integration. The enhanced power at the intermodulation frequency thus reflects the ease of lower-order audiovisual integration and demonstrates that speech-gesture information interacts in higher-order language areas. Furthermore, we provide a proof-of-principle of the use of RIFT to study the integration of audiovisual stimuli, in relation to, for instance, semantic context.

## Introduction

During communication in real-life settings, our brain needs to integrate auditory input with visual input in order to form a unified percept of the environment. Several magneto- and electroencephalography (M/EEG) studies have demonstrated that integration of non-semantic audiovisual inputs can occur as early as 50-100 ms after stimulus onset (e.g., Giard & Peronnet, 1999; Molholm et al., 2002; Talsma et al., 2010), and encompasses a widespread network of primary sensory and higher-order regions (e.g., Beauchamp et al., 2004; Calvert, 2001; Werner & Noppeney, 2010).

The integration of these audiovisual inputs has been studied using frequency tagging (Giani et al., 2012; Regan et al., 1995). Here, an auditory or visual stimulus is periodically modulated at a specific frequency, for example by modulating the luminance of a visual stimulus or the amplitude of an auditory stimulus. This produces steady-state evoked potentials (SSEPs, SSEFs for MEG) with strong power at the tagged frequency (for frequency-tagging in the visual domain and steady-state visual evoked responses (SSVEP), see e.g. Norcia et al., 2015; Vialatte et al., 2010; Gulbinaite et al., 2019, for frequency tagging in the auditory domain and auditory steady-state responses (ASSR), see e.g. Baltus & Herrmann, 2015; Picton et al., 2003; Ross et al., 2005; Ross et al., 2003). This technique is especially interesting in the context of studying audiovisual integration, because it enables the tagging of an auditory stimulus and a visual stimulus at two different frequencies (f_visual_ and f_auditory_) in order to study whether and how these two inputs interact in the brain. Previous work has suggested that when the auditory and visual signals interact, this results in increased power at the intermodulation frequencies of the two stimuli (e.g., |f_visual_ - f_auditory_| or f_visual_ + f_auditory_) (Regan & Regan, 1989). Such intermodulation frequencies arise from nonlinear interactions of the two oscillatory signals. In the case of audio-visual integration, the intermodulation likely reflects neuronal activity that combines the signals of the two inputs beyond linear summation (Regan & Regan, 1988; Zemon & Ratliff, 1984).

However, other authors have reported inconclusive results on the occurrence of such intermodulation frequencies as a signature of nonlinear audiovisual integration in neural signals. Furthermore, this integration has so far only been studied in non-semantic contexts (e.g., the integration of tones and gratings). For example, whereas Regan et al. (1995) identified intermodulation frequencies (i.e., as a result of tagging an auditory and visual stimulus) in an area close to the auditory cortex, Giani et al., (2012) identified intermodulation frequencies within (i.e., as a result of tagging two signals in the visual domain), but not between modalities (i.e., as a result of tagging both an auditory and a visual signal).

In both of these previous studies, frequency tagging was applied at relatively low frequencies (< 30 Hz for visual stimuli, < 40 Hz for auditory stimuli) (Giani et al., 2012; Regan et al., 1995). This might be problematic, considering that spontaneous neuronal oscillations at lower frequencies (e.g., alpha and beta oscillations) are likely entrained by frequency tagging (Keitel et al., 2014; Spaak et al., 2014). In the current study, we use novel projector technology to perform frequency tagging at high frequencies (rapid invisible frequency tagging; RIFT), and in a semantic context. Previous work has demonstrated that neuronal responses to a rapidly flickering LED can be driven and measured up to 100 Hz (Herrmann, 2001), and can successfully be used to study sensory processing in the brain (Herring, 2017; Zhigalov et al., 2019). We here leverage these rapid neural responses in order to circumvent the issue of endogenous rhythms interacting with low-frequency tagging signals.

We use speech-gesture integration as a test case for studying rapid invisible frequency tagging in a semantic context. Speech-gesture integration is a form of semantic audiovisual integration that often occurs in natural, face-to-face communication. Previous behavioral and neuroimaging studies have demonstrated that listeners process and integrate speech and gestures at a semantic level, and that this integration relies on a network involving left inferior frontal gyrus (LIFG), left-temporal regions (STS/MTG), motor cortex, and visual cortex (Dick et al., 2014; Drijvers, Ozyürek, et al., 2018; Drijvers, Ozyurek, et al., 2018; Drijvers et al., 2019; Holle et al., 2008, 2010; Kircher et al., 2009; Straube et al., 2012; Willems et al., 2007, 2009; Zhao et al., 2018). Using frequency tagging in such a context to study whether intermodulation frequencies can be identified as a signature of nonlinear audiovisual integration would provide a proof-of-principle for the use of such a technique to study the integration of multiple inputs during complex dynamic settings, such as multimodal language comprehension.

In the present study, we set out to explore whether RIFT can be used to identify intermodulation frequencies as a result of the interaction between a visual and auditory tagged signal in a semantic context. Participants watched videos of an actress uttering action verbs (tagged at f_auditory_ = 61 Hz) accompanied by a gesture (tagged at f_visual_ = 68 Hz). Integration difficulty of these inputs was modulated by auditory factors (clear/degraded speech) and visual factors (congruent/incongruent gesture). For the visually tagged input, we expected power to be strongest at 68 Hz in occipital regions. For the auditory tagged input, we expected power to be strongest at 61 Hz in auditory regions. We expected the interactions between the visually tagged and auditory tagged signal to be non-linear in nature, resulting in spectral peaks at the intermodulation frequencies of f_visual_ and f_auditory_ (i.e., f_visual_ + f_auditory_ and f_visual_ – f_auditory_). On the basis of previous work (e.g., Drijvers, Ozyurek & Jensen, 2018a/b, 2019), we expected the locus of the intermodulation frequencies to occur in LIFG and left-temporal regions such as pSTS/MTG, areas known to be involved in speech-gesture integration.

## Methods

### Participants

Twenty-nine right-handed native Dutch-speaking adults (age range = 19 - 40, mean age = 23.68, SD = 4.57, 18 female) took part in the experiment. All participants reported normal hearing, normal or corrected-to-normal vision, no neurophysiological disorders and no language disorders. All participants were recruited via the Max Planck Institute for Psycholinguistics participant database and the Radboud University participant database, and gave their informed consent preceding the experiment. Three participants (2 females) were excluded from the experiment due to unreported metal in dental work (1) or excessive motion artifacts (>75% of trials affected) (2). The final data set included the data of 26 participants.

### Stimulus materials

Participants were presented with 160 video clips showing an actress uttering a highly-frequent action verb accompanied by a matching or a mismatching iconic gesture (see for a detailed description of pre-tests on recognizability and iconicity of the gestures, (Drijvers & Ozyürek, 2017)). All gestures used in the videos were rated as potentially ambiguous when viewed without speech, which allowed for mutual disambiguation of speech and gesture (Habets et al., 2011).

In all videos, the actress was standing in front of a neutrally colored background, in neutrally colored clothes. We predefined the verbs that would form the ‘mismatching gesture’, in the sense that we asked the actress to utter the action verb, and depict the other verb in her gesture. This approach was chosen because we included the face and lips of the actress in the videos, and we did not want to recombine a mismatching audio track to a video to create the mismatch condition. Videos were on average 2000 ms long (SD = 21.3 ms). After 120 ms, the preparation (i.e., the first frame in which the hands of the actress moved) of the gesture started. On average, at 550 ms (SD = 74.4 ms), the meaningful part of the gesture (i.e., the stroke) started, followed by speech onset at 680 ms (SD = 112.54 ms), and average speech offset at 1435 ms (SD = 83.12 ms) None of these timings differed between conditions. None of the iconic gestures were prescripted. All gestures were performed by the actress on the fly.

All audio files were intensity-scaled to 70 dB and denoised using *Praat* (Boersma & Weenink, 2015), before they were recombined with their corresponding video files using Adobe Premiere Pro. For 80 of the 160 sound files, we created noise-vocoded versions using *Praat*. Noise-vocoding pertains the temporal envelope of the audio signal, but degrades the spectral content (Shannon et al., 1995). We used 6-band noise-vocoding, as we demonstrated in previous work that this is the noise-vocoding level where the auditory signal is reliable enough for listeners to still be able to use the gestural information for comprehension (Drijvers & Ozyürek, 2017). To achieve this, we band-pass filtered the sound files between 50 and 8000 Hz in 6 logarithmically spaced frequency bands with cut-off frequencies at 50, 116.5, 271.4, 632.5, 1473.6, 3433.5 and 8000 Hz. These frequencies were used to filter white noise and obtain six noise bands. We extracted the amplitude envelope of each band using half-wave rectification and multiplied the amplitude envelope with the noise bands. These bands were then recombined. Sound was presented to participants using MEG-compatible air tubes.

We manipulated integration strength in the videos by auditory (clear/degraded) and visual (congruent/incongruent) factors (see Figure 1). This resulted in four conditions: clear speech + matching gesture (CM), clear speech + mismatching gesture (CMM), degraded speech + matching gesture (DM) and degraded speech + mismatching gesture (DMM). These stimuli have been thoroughly pretested and used in previous work on speech-gesture integration (e.g., Drijvers & Ozyurek, 2017; Drijvers, Ozyurek & Jensen, 2018). All of the conditions contained 40 videos. All verbs and gestures were only presented once. Participants were asked to pay attention to the videos and identify what verb they heard in the videos in a 4-alternative forced choice identification task.

**Figure 1.**
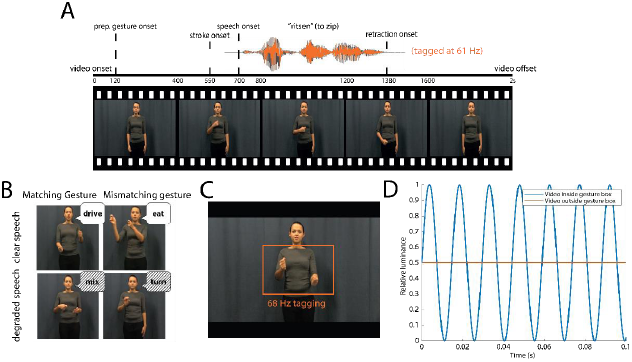
A. Illustration of the structure of the videos. Speech was amplitude-modulated at 61 Hz. B. Illustration of the different conditions. C. Area used for visual frequency tagging at 68 Hz. D. Illustration of luminance manipulation for visual-frequency tagging. D. Frequency tagging was achieved by multiplying the luminance of the pixels with a 68 Hz sinusoid. Modulation signal was equal to 0.5 at sine wave zero-crossing to preserve the mean luminance of the video, and was phase-locked across trials.

### Procedure

Participants were tested in a dimly-lit magnetically shielded room and seated 70 cm from the projection screen. All stimuli were presented using MATLAB 2016b (Mathworks Inc, Natrick, USA) and the Psychophysics Toolbox, version 3.0.11 (Brainard, 1997; Kleiner et al., 2007; Pelli, 1997). To achieve rapid invisible frequency tagging, we used a GeForce GTX960 2GB graphics card with a refresh rate of 120 Hz, in combination with a PROPixx DLP LED projector (VPixx Technologies Inc., Saint-Bruno-de-Montarville, Canada), which can achieve a presentation rate up to 1440 Hz. This high presentation rate is achieved by the projector interpreting the four quadrants and three colour channels of the GPU screen buffer as individual smaller, grayscale frames, which it then projects in rapid succession, leading to an increase of a factor 12 (4 quadrants * 3 colour channels * 120 Hz = 1440 Hz) (User Manual for ProPixx, VPixx Technologies Inc., Saint-Bruno-de-Montarville, Canada).

#### Frequency tagging

The area of the video that would be frequency-tagged was defined by the rectangle in which all gestures occurred, which measured 10.0 by 6.5 degrees of visual angle (width by height). The pixels within that area were always tagged at 68 Hz. This was achieved by multiplying the luminance of the pixels within that square with a 68 Hz sinusoid (modulation depth = 100 %; modulation signal equal to 0.5 at sine wave zero-crossing, in order to preserve the mean luminance of the video), phase-locked across trials (see Figure 1D). For the auditory stimuli, frequency tagging was achieved by multiplying the amplitude of the signal with a 61 Hz sinusoid, with a modulation depth of 100 % (following (Lamminmäki et al., 2014)). In a pretest, we presented 11 native Dutch speakers with half of the stimuli containing the amplitude modulation, and half of the stimuli not containing the amplitude modulation in both clear and degraded speech. Participants were still able to correctly identify the amplitude modulated stimuli in clear speech (mean % correct without amplitude modulation: 99.54, with amplitude modulation: 99.31) and in degraded speech (mean % correct without amplitude modulation: 72.74, with amplitude modulation: 70.23) and did not suffer more compared to when the signal was not amplitude modulated.

Participants were asked to attentively watch and listen to the videos. Every trial started with a fixation cross (1000 ms), followed by the video (2000 ms), a short delay period (1500 ms), and a 4-alternative forced choice identification task (max 3000 ms, followed by the fixation cross of the next trial as soon as a participant pressed one of the 4 buttons). In the 4-alternative forced choice identification task, participants were presented with four written options, and had to identify which verb they heard in the video by pressing one of 4 buttons on an MEG-compatible button box. This task ensured that participants were attentively watching the videos, and to check whether the verbs were understood. Participants were instructed not to blink during video presentation.

Throughout the experiment, we presented all screens at a 1440 Hz presentation rate. Brain activity was measured using MEG, and was recorded throughout the experiment. The stimuli were presented in four blocks of 40 trials each. The whole experiment lasted approximately 30 minutes, and participants were allowed to take a self-paced break after every block. All stimuli were presented in a randomized order per participant.

### MEG data acquisition

MEG was recorded using a 275-channel axial gradiometer CTF MEG system (CTF MEG systems, Coquitlam, Canada). We used an online low-pass filter at 300 Hz and digitized the data at 1200 Hz. All participants’ eye gaze was recorded by an SR Research Eyelink 1000 eye tracker for artifact rejection purposes. The head position of the participants was tracked in real time by recording markers on the nasion, and left and right periauricular points (Stolk et al., 2013). This enabled us to readjust the head position of participants relative to their original starting position whenever the deviation was larger than 5 mm. After the experiment, T1-weighted structural magnetic resonance images (MRI) were collected from 24 out of 26 participants using a Siemens 3T MAGNETOM Skyra system.

### MEG data analysis

#### Preprocessing

All MEG data were analyzed using the FieldTrip toolbox (version 20180221) (Oostenveld et al., 2011) running in a Matlab environment (2017b). All data were segmented into trials starting 1 s before and ending 3 s after the onset of the video. The data were demeaned and line noise was attenuated using a discrete Fourier transform approach at 50, 100 and 150 Hz. All trials that contained jump artifacts or muscle artifacts were rejected using a semi-automatic routine. The data were then down-sampled to 400 Hz. Independent component analysis (Bell & Sejnowski, 1995; Jung et al., 2000) was used to remove residual eye movements and cardiac-related activity (average number of components removed: 6.05). All data were then inspected on a trial-by-trial basis to remove artifacts that were not identified using these rejection procedures. These procedures resulted in rejection of 8.3 % of the trials. The number of rejected trials did not differ significantly between conditions.

#### Frequency tagging analyses - Sensor-level

To investigate the response in auditory and visual regions to the frequency-tagged signal, we first calculated event-related fields by averaging time-locked gradiometer data over trials, over conditions, and over participants. All tagged stimuli were presented phase-locked over trials. We used an approximation of planar gradiometer data to facilitate interpretation of the MEG data, as planar gradient maxima are thought to be located above the neuronal sources that may underlie them (Bastiaansen & Knösche, 2000). This was achieved by converting the axial gradiometer data to orthogonal planar gradiometer pairs, which were combined by using root-mean-square (RMS) for the ERFs. For the power analyses, we computed the power separately for the two planar gradient directions, and combined the power data by averaging the two. To visualize the responses per tagging frequency (Figure 3), we used a notch (i.e. band-stop) filter between 60 and 62 Hz to display the ERF at 68 Hz, and a notch filter between 67 and 69 Hz to display the ERF at 61 Hz.

We then performed a spectral analysis on an individual’s ERF data pooled over conditions, in the time window in which both the auditory and visual stimulus unfolded (0.5 - 1.5 s), and a post-stimulus baseline (2.0 - 3.0s). We chose this post-stimulus time window as a baseline because, contrary to the pre-stimulus time window, it is not affected by the button press of the 4-alternative forced choice identification task. We chose the 0.5-1.5 s time window to focus our analysis on, because this time window captures both the meaningful part of the gesture and the full speech signal. We computed power spectra in frequencies ranging from 1 to 130 Hz for both the baseline and stimulus window using fast Fourier transform and a single Hanning taper of the 1s segments. This data was then averaged over conditions, and the stimulus window was compared to the baseline window.

#### Frequency tagging analyses - Source-level

To reconstruct activity at the tagging frequencies, we calculated coherence between a pure sine wave at either 61 Hz or 68 Hz, reflecting the tagged stimulus, and the observed MEG signal at those frequencies. Although the phase of the tagging was designed to be identical over trials, the projector that we used occasionally experienced a brief delay in presenting the video material (in 16 of the 26 participants). We corrected for this by translating any observed delays between video onset and offset markers (recorded in a stimulus trigger channel) into a phase-difference, which was then subtracted from the tagging signal. Note that this correction only uses information in the stimulus marker channel and the length of the original video files, and does not rely on any information in the measured MEG signal.

We performed source analysis to identify the neuronal sources that were coherent with the modulation signal at either 61 Hz or 68 Hz, and compared the difference in coherence in the stimulus and post-stimulus window. This was done pooled over conditions. Source analyses on coherence values (for 61 and 68 Hz) and power values (for the intermodulation frequency at 7 Hz, see results), was performed using dynamic imaging of coherent sources (DICS; (Gross et al., 2001)) as a beamforming approach. We computed a common spatial filter per subject from the lead field matrix and the cross-spectral density matrix (CSD) that was the same for all conditions. An individual’s leadfield was obtained by spatially co-registering an individual’s anatomical MRI to the MEG data by the anatomical markers at the nasion and left and right periaucular points. Then, for each participant, a single-shell head model was constructed on the basis of the MRI (Nolte, 2003). A source model was created for each participant by warping a 10 mm spaced grid defined in MNI space to the individual participant’s segmented MRI. The MNI template brain was used for those participants (2/26) for which an individual MRI scan was not available.

After establishing regions that showed elevated coherence with the tagged stimuli, we proceeded to test the effect of the experimental conditions (clear versus degraded speech; matching versus mismatching gesture) within these regions-of-interest (ROIs). The ROIs for the auditory and visual tagged signals were defined by taking the grid points that exceeded 80 percent of the peak coherence difference value between stimulus and baseline, across all conditions. For these ROIs, coherence difference values were extracted per condition. Analogously, the ROI for the intermodulation frequency at 7 Hz was defined by taking those grid points that exceeded 80 percent of the peak power difference value between stimulus and baseline. The 80 percent threshold was chosen as an exploratory threshold.

#### Statistical comparisons

As we predefined our frequencies of interest and have specific regions of interest for analysis, we compared the differences between conditions using 2×2 repeated measures ANOVAs, with the factors Speech (clear/degraded) and Gesture (matching/mismatching).

## Results

Participants watched videos of an actress uttering action verbs in clear or degraded speech, accompanied by a matching or mismatching gesture. After the video, participants were asked to identify the verb they heard in a 4-alternative forced choice identification task, presented on the screen in written form. Video presentation was manipulated by tagging the gesture space in the video by 68 Hz flicker, while the sound in the videos was tagged by 61 Hz amplitude modulation (see Figure 1).

### Behavioral results

In our behavioral task we replicated previous results (see Drijvers, Ozyürek, et al., 2018; Drijvers & Özyürek, 2018) and observed that when the speech signal was clear, response accuracy was higher than when speech was degraded (F(1, 25) = 301.60, *p* < .001, partial η^2^ = .92) (mean scores and SDs: CM: 94.7% (SD = 4.0%), CMM: 90.2% (SD = 5.6%), DM: 85.0% (SD = 8.2%), DMM: 66.5% (SD = 7.8%)). Similarly, response accuracy was higher when a gesture matched compared to mismatched the speech signal (F(1, 25) = 184.29, *p* < .001, partial η^2^ = .88). The difference in response accuracy was larger in degraded speech than in clear speech (F(1, 25) = 4.87, *p* < .001, partial η^2^ = .66) (see raincloud plots (Allen et al., 2019), Figure 2).

**Figure 2.**
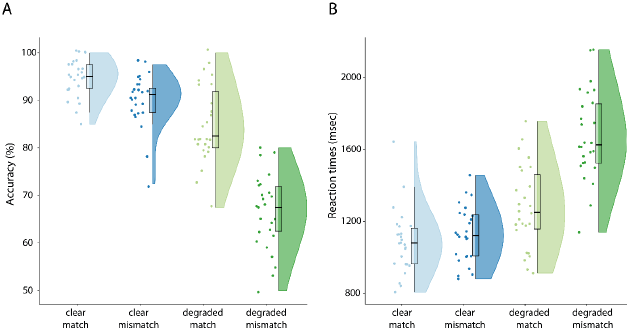
A: Accuracy results per condition. Response accuracy is highest for clear speech conditions, and when a gesture matches the speech signal. B: Reaction times per condition. Reaction times are faster in clear speech and when a gesture matches the speech signal. Raincloud plots reveal raw data, density and boxplots for coherence change.

We observed similar results in the reaction times (RTs). Participants were faster to identify the verbs when speech was clear, compared to when speech was degraded (F(1, 25) = 198,06, *p* < .001, partial η^2^ = .89) (mean RTs and SDs: CM: 1086.3 ms, SD = 177.1 ms, CMM: 1127.92 ms, SD = 153.84 ms, DM: 1276.96 ms, SD = 230.13 ms, DMM: 1675.77 ms, SD = 246.69 ms). Participants were faster to identify the verbs when the gesture matched the speech signal, compared to when the gesture mismatched the speech signal (F(1, 25) = 105,42, *p* < .001, partial η^2^ = .81). This difference in reaction times was larger in degraded speech than in clear speech (F(1, 25) = 187,78, *p* < .001, partial η^2^ = .88).

In sum, these results demonstrate that gestures facilitate speech comprehension when the actress performed a matching gesture, but hindered comprehension when she performed a mismatching gesture. This effect was larger in degraded speech than in clear speech.

### MEG results - Frequency tagging

#### Both visual and auditory frequency tagging produce a clear steady-state response that is larger than baseline

As a first step, we calculated the time-locked averages of the event-related fields pooled over conditions. Auditory frequency tagging at 61 Hz produced an auditory steady-state response over left and right-temporal regions (see Figure 3A), and visual frequency tagging at 68 Hz produced a clear visual steady-state response at occipital regions (see Figure 3B).

**Figure 3:**
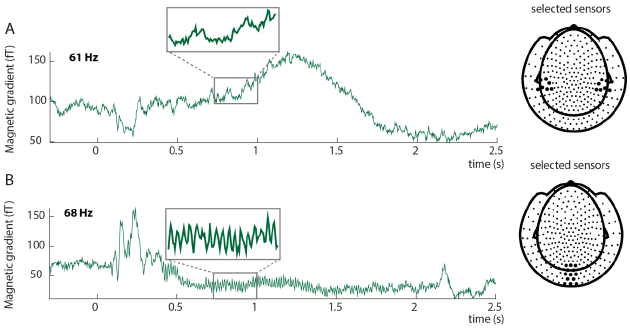
Event-related fields show clear responses at the tagged frequencies. Auditory input was tagged by 61 Hz amplitude modulation (A), Visual input was tagged by 68 Hz flicker (B). The insets reflect an enlarged part of the signal to clearly demonstrate the effect of the tagging on the event-related fields. Tagging was phase-locked over trials. A: Average ERF for a single subject at selected sensors overlying the left and right temporal lobe. The highlighted sensors in the right plot reflect the sensors for which the ERF is plotted. B: Average ERF for 68 Hz for a single subject at selected sensors overlying occipital cortex. The highlighted locations in the right plot reflect the sensors for which the ERF is plotted. ERFs show combined planar gradient data.

To explicitly compare the tagged signals between stimulus (0.5 – 1.5 s) and post-stimulus baseline (2.0 – 3.0 s) periods, we plotted the difference in spectral power calculated from the ERF (i.e. power of the time-locked average) in Figure 4. We observe that both visual and auditory responses at the tagged frequency were reliable larger in the stimulus period than in the baseline (see below for statistical assessment at the source level). Note that the visual tagged signal at 68 Hz seems to be more focal and strong than the auditory tagged signal at 61 Hz (see Figure 4). These analyses confirm that we were able to induce high-frequency steady-state responses simultaneously for both auditory and visual stimulation.

**Figure 4:**
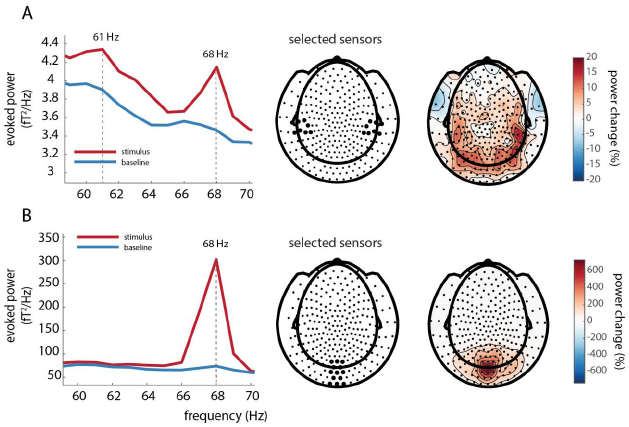
A: Power over auditory sensors peaks at the tagged frequency of the auditory stimulus (61 Hz). Note the visual 68 Hz tagged signal is still observable at left- and right-temporal sensors of interest. 61 Hz power is stronger in the stimulus interval than in the baseline interval, and is widely spread over posterior regions, with maxima at right-temporal regions. B: A power increase is observed at the tagged frequency (68 Hz) for the visual stimuli. 68 Hz power is larger in the stimulus than in the baseline window and is strongest over occipital regions.

#### Coherence is strongest at occipital regions for the visually tagged signal (68 Hz) and strongest when speech is clear

We proceeded to identify the neural generators of the tagged signals using beamformer source analysis. We computed source-level coherence coefficients for all conditions pooled together. This was done by computing coherence between a visual dummy 68 Hz modulation signal and the observed MEG data. The relative coherence increase between stimulus and baseline was largest in occipital regions (see Figure 5A), in an area consistent with early visual cortex.

**Figure 5:**
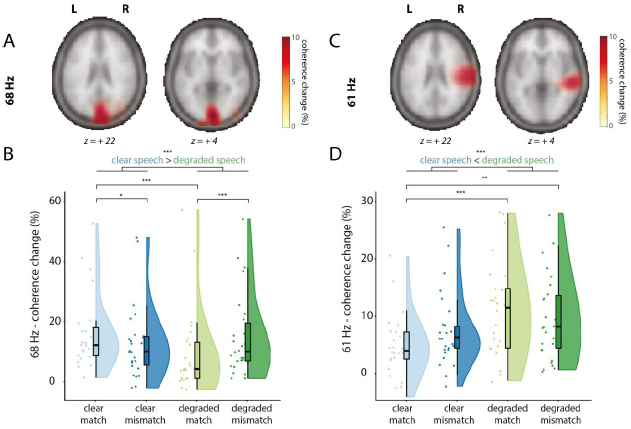
Sources of the visually tagged signal at 68 Hz (A/B) and sources of the auditory tagged signal at 61 Hz (C/D), and individual scores in the respective ROI per condition (clear match/clear mismatch/degraded match/degraded mismatch. Z-coordinates of slices are in mm and in MNI space. A: Coherence change in percentage when comparing coherence values in the stimulus window to a post-stimulus baseline for 68 Hz (the frequency of the visual tagging), pooled over conditions. Only positive coherence change values are plotted (>80% of peak maximum). Coherence change is largest over occipital regions for the visually tagged signal. B: Coherence change values in percentage extracted from the 68 Hz ROI. Raincloud plots reveal raw data, density and boxplots for coherence change. C: Coherence change in percentage when comparing coherence values in the stimulus window to a post-stimulus baseline for 61 Hz (the frequency of the auditory tagging), pooled over conditions. Only positive coherence values are plotted (>80% of peak maximum). Coherence change is largest over right-temporal regions. D: Coherence change values in percentage extracted from the 61 Hz ROI. Raincloud plots reveal raw data, density and boxplots for coherence change.

To compare conditions, we then formed a visual ROI by selecting those grid points exceeding an exploratory threshold of 80 % of the peak coherence increase. For each participant, the percentage of change in coherence between stimulus and baseline was computed in that ROI per condition and compared in a 2×2 (Speech: clear/degraded, Gesture: matching/mismatching) RM-ANOVA (see Figure 5B). Coherence change was larger for videos containing clear speech than videos containing degraded speech (F(1, 25) = 17.14, *p* < .001, partial η2 = .41), but did not differ between matching or mismatching trials (F(1, 25) = 0.025, *p* = .87, partial η2 = .001). We observed a significant interaction between Speech and Gesture (F(1, 25) = 26.87, *p* < .001, partial η2 = .52). Post-hoc pairwise comparisons revealed a stronger coherence change in videos containing clear speech and a matching gesture (CM) than clear speech and a mismatching gesture (CMM) (t(25) = 3.26, *p* = .015), and a stronger coherence change in videos containing degraded speech and a mismatching gesture (DMM) than in videos containing degraded speech and a matching gesture (DM) (t(25) = -4.03, *p* < .001). Coherence change was larger in CM than in DM (t(25) = 6.59, *p* < .001), in CMM than DM (t(25) = 2.93, *p* = .04), but not larger in CM than in DMM (t(25) = 2.02, *p* = .27), and not larger in CMM compared to DMM (t(26) = −1.74, *p* = .48).

These results thus indicate that visual regions responded stronger to the frequency-tagged gestural signal when speech was clear than when speech was degraded. This suggests that when speech is clear, participants allocate more visual attention to gestures than when speech is degraded, especially when a gesture matched the speech signal. When speech is degraded, participants allocate more attention to mismatching than to matching gestures.

#### Coherence is strongest at right-temporal regions for the auditory tagged signal (61 Hz) and strongest when speech is degraded

Similar to the visually tagged signal, we first computed coherence coefficients for all conditions pooled together. This was done by computing source-level coherence between a dummy 61 Hz modulation signal (reflecting the auditory tagging drive) and the observed MEG data. The coherence difference between stimulus and baseline peaked at right temporal regions (Figure 5C), in an area consistent with (right) early auditory cortex.

To compare conditions, we then formed the auditory ROI by selecting those grid points exceeding an exploratory threshold of 80 % of peak coherence change. Again, coherence change values per condition and per participant were compared in a 2×2 RM-ANOVA (see Figure 5D). Coherence change was larger in degraded speech conditions than in clear speech conditions (F(1, 25) = 12.87, *p* = .001, partial η2 = .34), but did not differ between mismatching and matching conditions (F(1, 25) = 0.09, *p* = .77, partial η2 = .04). No interaction effect was observed (F(1, 25) = 3.13, *p* = .089, partial η2 = .11). Post-hoc pairwise comparisons revealed that there was no difference in coherence change when comparing CM and CMM (t(25) = -1.44, *p* = .81), or between DM and DMM (t(25) = 1.38, *p* = .90). Coherence change was larger in DM than in CM (t(25) = -4.24, *p* < .001), and in DMM than in CM (t(25) = -3.90, *p* < .01) but not when comparing CMM to DMM (t(25) = -1.40, *p* = .87). These results thus indicate that right-lateralized auditory regions processed the frequency-tagged auditory signal more strongly when speech was degraded than when speech was clear. This suggests that when speech is degraded, participants allocate more auditory attention to speech than when speech is clear.

#### An intermodulation frequency was observed at 7 Hz (|f_visual_ - f_auditory_|), but not at 129 Hz (f_visual_ + f_auditory_)

To test whether intermodulation frequencies (|f_visual_ - f_auditory_|, f_visual_ + f_auditory_) could be observed, we then calculated power spectra of the ERFs in the stimulus time window and the post-stimulus time window at 7 Hz and 129 Hz. Only for 7 Hz a difference between stimulus and baseline was observed at left frontal and left temporal sensors (Figure 6A/C). No reliable differences were observed for 129 Hz (Figure 6D). Interestingly, the spectral peak at 7 Hz during stimulus was most pronounced for the clear/match condition (Figure 6E).

**Figure 6:**
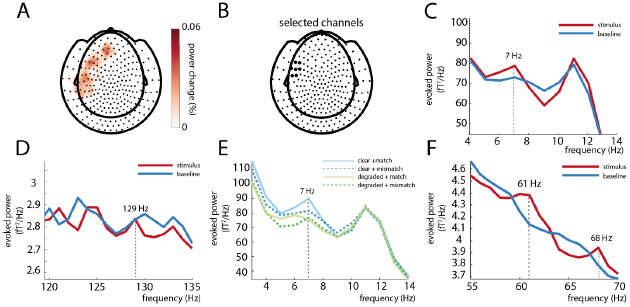
An intermodulation frequency could be observed at 7 Hz (|f_visual_-f_auditory_|) (A/C/E) but not 129 Hz (f_visual_+f_auditory_). (D). A: 7 Hz power in the stimulus window is larger than baseline over left-temporal and left-frontal sensors. Only positive values are plotted. B: Selected sensors (based on visual inspection). The black highlighted sensors represent the sensors at which the power spectra of the ERFs was calculated. C: Power spectra of 7 Hz (stimulus>baseline). D: No difference could be observed at 129 Hz between stimulus and baseline. E: Power spectra per condition. 7 Hz power peaks strongest in the clear+match condition. F: Power spectra of 61 Hz and 68 Hz over selected channels of 7 Hz power peak (see B).

As a next step, we then took a similar approach as for the visual and auditory tagged stimuli and calculated the coherence difference between stimulus and baseline at 7 Hz, pooled over conditions. This was done by computing source-level coherence between a dummy 7 Hz modulation signal (the intermodulation frequency of our 61 and 68 Hz tagging signals, specified as the multiplication of the 61 and 68 Hz dummy signal) and the observed MEG data. The coherence analysis did not reveal any differences between stimulus and baseline (see Figure 7A). It should be noted here that our frequency-tagged signals at f_auditory_ and f_visual_ were exactly phase-consistent across trials, since the phase was uniquely determined by the stimuli themselves. However, it is possible that the phase of the intermodulation signal has a much weaker phase consistency across trials, since it depends not only on the stimuli but also on the nature of the nonlinear neural interaction. If this is the case, we might still observe an effect on the *power* at the intermodulation frequency, rather than the coherence. We therefore performed source analysis on the power of the combined conditions versus baseline. Here, we observed a power change at 7 Hz in left frontal and temporal regions that mirrored the effect we observed at sensor level (Figure 7B).

**Figure 7:**
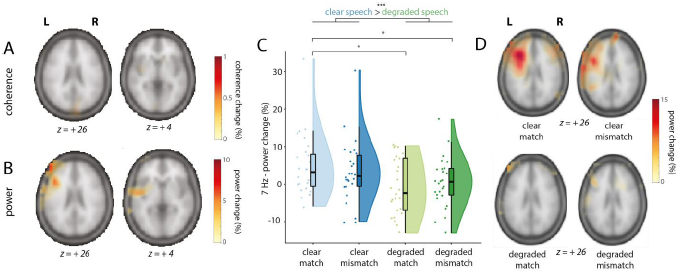
Sources of the intermodulation frequency (f_visual_-f_auditory_) at 7 Hz and individual scores in the left-frontotemporal ROI per condition (clear match/clear mismatch/degraded match/degraded mismatch). Z-coordinates of slices are in mm and in MNI space. A: Coherence change in percentage when comparing coherence values in the stimulus window to a post-stimulus baseline for 7 Hz (intermodulation frequency, f_visual_ - f_auditory_), pooled over conditions. Only positive coherence values are plotted (> 80 % of maximum). No differences could be observed. B: Power change in percentage when comparing power values in the stimulus window to a post-stimulus baseline for 7 Hz, pooled over conditions. Power changes were largest in left-frontal and left-temporal regions. Highest peak value was at MNI coordinates -44, 24, 22, and extended from LIFG to pSTS/MTG. Only positive coherence values are plotted (> 80 % of maximum). C: Power change values in percentage extracted from the 7 Hz ROI in source space. Raincloud plots reveal raw data, density and boxplots for power change per condition. D: Power change in percentage when comparing power values in the stimulus window to a post-stimulus baseline for 7Hz, per condition.

The condition-averaged effect at the intermodulation frequency of 7 Hz is less striking than at the primary tagged frequencies of 61 and 68 Hz, potentially due to it being driven mainly by one of the four conditions only (see Figure 6E). Note that the 61 and 68 Hz signal were still present over the left-frontotemporal sensors where we observed the 7Hz effect (see Figure 6F). As a next step, and sticking to our a priori defined hypotheses and analysis plan, we again proceeded by comparing conditions within an ROI defined by the condition-averaged contrast in source space. As before, the ROI was defined as those grid points exceeding an exploratory threshold of 80 % of the peak power change from baseline to stimulus epochs. We compared the strength of the 7 Hz signal at source level between conditions by using a 2×2 RM-ANOVA (Figure 7C). Power change was larger in clear speech conditions than in degraded speech conditions (F(1, 25) = 10.26, *p* = .004, partial η2 = .29), but did not differ between matching and mismatching trials (F(1, 25) = 0.01, *p* = .91, partial η2 = .001), suggesting an effect of speech degradation, but not of semantic congruency. No interaction effect was observed (F(1, 25) = 1.27, *p* = .27, partial η2 = .05). Post-hoc pairwise comparisons revealed that 7 Hz power was not different for CM compared to CMM (t(25) = 1.14, *p* = 1), and not different for DM compared to DMM (t(25) = -.67, *p* = 1). However, 7 Hz power was larger in CM than in DM (t(25) = 3.01, *p* = .025), and larger in CM than in DMM (t(25) = 2.82, *p* = .045). No difference was observed between CMM and DMM (t(25) = 1.61, *p* = .6). To rule out that these differences in 7 Hz power were due to general power differences in the theta band, we compared the strength of 6 Hz and 8 Hz between conditions, using two 2×2 RM-ANOVA’s. Here, no differences between conditions were observed (all *p* > 0.05), suggesting this was specific to the 7 Hz signal. These results are also in line with previous MEG studies on speech-gesture integration, where no differences in theta power were observed (Drijvers, Ozyürek, et al., 2018; Drijvers, Ozyurek, et al., 2018b; Drijvers, van der Plas, et al., 2019).

In addition to our ROI-based analysis, we present the full beamformer source maps of 7 Hz power (stimulus versus baseline) for the four conditions in Figure 7D. These reveal results fully compatible with the aforementioned RM-ANOVA. Furthermore, they show that our ROI selection on the condition-averaged response versus baseline was likely suboptimal, since the source map for CM shows a more clearly elevated intermodulation cluster than the average (in line with the sensor-level results shown in Figure 6A).

These results thus demonstrate that we could reliably observe an intermodulation signal when speech was clear and a gesture matched the speech signal. Left-frontotemporal regions showed a stronger intermodulation peak (reflecting the lower-order interaction between the auditory and visually tagged signal) when speech was clear than when speech was degraded. This suggests that the interaction between the auditory and visual tagged signal is strongest when signal quality was optimal and speech was clear.

## Discussion

In the current MEG study we provide a proof-of-principle that rapid invisible frequency tagging (RIFT) can be used to estimate task-dependent neuronal excitability in visual and auditory areas, as well as the auditory-visual interaction. Coherence was strongest over occipital regions for the visual-tagged input, and strongest when speech was clear. Coherence was strongest over right-temporal regions for the auditory-tagged input and strongest when speech was degraded. Importantly, we identified an intermodulation frequency at 7 Hz (f_visual_ - f_auditory_) as a result of the interaction between a visual frequency-tagged signal (gesture; 68 Hz) and an auditory frequency-tagged signal (speech; 61 Hz). In line with our hypotheses, power at this intermodulation frequency was strongest in LIFG and left-temporal regions (pSTS/MTG), and was strongest when the lower-order integration of auditory and visual information was optimal (i.e., when speech was clear). Below we provide interpretations of these results.

### Clear speech enhances visual attention to gestural information

In occipital regions, we observed a stronger drive by the 68 Hz visual modulation signal when speech was clear than when speech was degraded. We speculate that this effect reflects that listeners allocate more visual attention to gestures when speech is clear. This speculative interpretation is in line with previous eye-tracking work that demonstrated that when speech is degraded, listeners gaze more often to the face and mouth than to gestures to extract phonological information to aid comprehension (Drijvers, Vaitonytė, et al., 2019), as well as previous work that revealed that the amplitude of SSVEPs was enhanced by visual attention, irrespective of whether the stimuli were task-relevant (Morgan et al., 1996; Müller et al., 2006). Note that gestural information is often processed in the periphery of a listener’s visual field (Gullberg & Holmqvist, 1999, 2002, 2006; Gullberg & Kita, 2009). As listeners do not necessarily need to extract the phonological information conveyed by the lips when speech is clear, overt visual attention might be directed to a ‘resting’ position in the middle of the screen during clear speech processing, resulting in stronger coherence with the visual drive when speech is clear than when speech is degraded. Pairwise comparisons of the conditions revealed that in clear speech, coherence was larger when the gesture matched, rather than mismatched, the signal. In line with the interpretation above, a listener might have reconsidered the auditory input when noticing that the gesture mismatched the perceived auditory input, and might have directed their attention to the face/lips of the actress, which, in turn, reduces visual attention to the gesture.

However, we observed the opposite effect when speech was degraded; i.e. a stronger coherence when the gesture mismatched, rather than matched, the degraded speech signal. We speculate that when speech is degraded and a gesture matches the signal, a listener might more strongly allocate visual attention to the information conveyed by the face/lips, so that information conveyed by the lips and the information conveyed by the gesture can jointly aid in disambiguating the degraded speech signal (Drijvers & Ozyürek, 2017). However, when speech is degraded and a gesture mismatches the signal, the uncertainty of both inputs may result in a reconsideration of both inputs, and thus a less fixed locus of attention (see also Nath & Beauchamp, 2011 for work on perceptual reliability weighting in clear and degraded speech). These interpretations are rather speculative, and further work is needed to disambiguate different interpretations. For example, future work could consider tagging the mouth-region to further investigate how a listener allocates visual attention to these two visual articulators during comprehension

### Degraded speech enhances auditory attention to speech information

In line with our hypotheses, we observed stronger drive by the 61 Hz amplitude modulation signal in temporal areas overlapping with auditory cortex when speech was degraded than when speech was clear. This response was strongest at right-temporal regions, which is in line with previous work that demonstrated that for speech stimuli, the ASSR is often localized to right-lateralized sources (Lamminmäki et al., 2014; Ross et al., 2005). Although both left- and right-hemispheres process speech, a right-lateralized dominance is often observed because right-lateralized regions are sensitive to spectral changes and prosodic information, and processing of low-level auditory cues (Zatorre & Gandour, 2008; Scott et al., 2000).

Previous work has reported enhanced ASSR responses to amplitude-modulated multi-speech babble when attention to this input increases (Keitel et al., 2011; Ross et al., 2004; Saupe et al., 2009; Talsma et al., 2010; Tiitinen et al., 1993). The enhanced ASSR which we observed in the degraded compared to clear speech conditions could thus reflect an increase in attention to the speech signal when speech is degraded. Note that no differences in coherence were observed when comparing matching and mismatching gestures in either clear or degraded speech. As the gesture congruency manipulation is a visual manipulation, this indicates that modulation of the ASSR is modality-specific (Parks et al., 2011; Rees et al., 2001).

### The auditory tagged speech signal and visual tagged gesture signal interact in left-frontotemporal regions

We set out to study whether intermodulation frequencies could be identified in a multimodal, semantic context as a result of the interaction of the visual and auditory tagged signals. In contrast to previous work by (Giani et al., 2012) using lower frequencies, we did observe an intermodulation frequency at 7 Hz (f_visual_ - f_auditory_), but not at 129 Hz (f_visual_ + f_auditory_). As responses in lower frequencies tend to be stronger than in higher frequencies, the higher-frequency intermodulation frequency might not have been identifiable as neurons cannot be driven in this fast range.

Note that although we observed a stronger 7 Hz power peak at sensor level in the stimulus interval compared to the baseline interval, we did not observe stronger coherence between a 7 Hz dummy signal and the observed MEG data at source level. This indicates that the phase of the intermodulation signal is not as consistent over trials as the f_visual_ and f_auditory_ signals, which in turn might imply that the time point of interaction of the two signals differs across trials. This could explain why we observed a clear difference between stimulus and baseline when we reconstructed the sources of the intermodulation frequency on the basis of power, but not coherence.

We observed a reliable peak at 7 Hz power during stimulation when integration of the lower-order auditory and visual input was optimal, i.e., when speech was clear. In line with our hypotheses, the source of the intermodulation frequency was localized in LIFG and left-temporal (pSTS/MTG) regions. It has been shown that these areas are involved in the integration of speech and gestures (Dick et al., 2014; Drijvers, Ozyürek, et al., 2018; Drijvers, Ozyurek, et al., 2018; Drijvers, van der Plas, et al., 2019; Holle et al., 2008, 2010; Kircher et al., 2009; Straube et al., 2012; Willems et al., 2007, 2009; Zhao et al., 2018). There are, however, important differences between the interpretation of the intermodulation frequency in this work, and the results observed in response to higher-order speech-gesture integration in previous work.

First, although previous work has observed effects related to higher-order integration in LIFG and pSTS/MTG, the observed intermodulation frequency in the current work is most likely related to lower-order integration. Specifically, we observed that power at the intermodulation frequency was stronger in clear speech conditions than in degraded speech conditions, but we did not observe an effect of gesture congruency. We therefore propose that, contrary to our hypotheses, power at the intermodulation frequency does not reflect the integration of higher-order semantic audiovisual integration, but rather is a direct reflection of the non-linear integration of lower-order speech and gesture information. This difference might be explained by considering that the intermodulation frequency is unable to capture higher-order effects that result from lexical access on the basis of the auditory and visual input. Second, the current work is not able to dissociate between the different roles of the LIFG and pSTS/MTG in the speech-gesture integration process. The accuracy of source modeling using MEG should be considered in the light of the inverse problem (Baillet, 2017). This limits our ability to make precise claims about the exact locus of the observed effect when comparing to fMRI (see e.g., Papeo et al., 2019, for a functional distinction of different subregions of the MTG in the speech-gesture integration process). Furthermore, fMRI is sensitive to modulations in the BOLD signal whereas MEG detects changes in neuronal synchronization. As such, these techniques provide complementary but not necessarily overlapping information on neuronal activation.

### Proof of principle: using RIFT to study the integration of complex and dynamic audiovisual stimuli in a semantic context

The current MEG study provides a proof of principle of the use of rapid invisible frequency tagging (RIFT) to study the integration of audiovisual stimuli, and is the first study to identify intermodulation frequencies as a result of the lower-order interaction between auditory and visual stimuli in a semantic context. Note that although previous work has reported the occurrence of intermodulation frequencies in a non-semantic context (Regan et al., 1995), other studies have failed to identify between-modality intermodulation frequencies (Giani et al., 2012). This could be due to the fact that lower frequencies were used for tagging. Another possibility is that this was due to the nature of the stimuli used in these studies. As Giani et al., (2012) suggest, the occurrence of intermodulation frequencies resulting from audiovisual integration of non-semantic inputs such as tones and gratings might reflect low-level spatiotemporal coincidence detection that is prominent for transient stimuli, but less so for sustained steady-state responses. Similarly, previous fMRI work that investigated the difference between transient and sustained BOLD responses revealed that primary auditory and visual regions were only involved in the integration of rapid transient stimuli at stimulus onset. However, integration for sustained responses did involve higher-order areas (Werner & Noppeney, 2011). The observed 7 Hz intermodulation frequency in response to our semantic audiovisual stimuli was also localized to higher-order areas, rather than early sensory regions. This again underlines the possibility that the observed intermodulation frequency in the current study reflects the ease of lower-order integration of these audiovisual stimuli in certain higher-order regions.

An important advantage of using RIFT is that spontaneous neuronal oscillations in lower frequencies were not entrained by our tagging frequencies. This might explain why a clear intermodulation frequency was observed in the current study, but was less easy to identify in previous work. Future studies might consider exploiting this feature and using RIFT to study the interaction of these endogenous lower frequency oscillations with the tagged signals, in order to elucidate their role in sensory processing. However, future work should also consider that high-frequency tagging might entrain spontaneous neuronal oscillations at higher frequencies. Although this was not directly relevant for the identification of the intermodulation frequency in this study, and we did not observe any gamma band modulations in response to the stimuli used in this study in earlier work (Drijvers, Ozyurek & Jensen, 2018b), it should be noted that gamma band modulations have been observed in other work related to linguistic semantic processing (e.g., in the 30-50 Hz range in Mellem et al., 2013; Wang et al., 2018).

### Conclusion

First of all, we provided a proof of principle that RIFT can be used to tag visual and auditory inputs at high frequencies, resulting in clear spectral peaks in the MEG signal, localized to early sensory cortices. Second, we demonstrated that RIFT can be used to identify intermodulation frequencies in a multimodal, semantic context. The observed intermodulation frequency was the result of the nonlinear interaction between visual and auditory tagged stimuli. Third, the intermodulation signal was localized to LIFG and pSTS/MTG, areas known to be involved in speech-gesture integration. The strength of this intermodulation frequency was strongest when lower-order signal quality was optimal. In conclusion, we thus propose that the strength of this intermodulation frequency reflects the ease of lower-order audiovisual integration, that RIFT can be used to study both unimodal sensory signals as well as their multimodal interaction in downstream higher-order areas, and that RIFT has many use cases for future work.

## Acknowledgements

This work was supported by Gravitation Grant 024.001.006 of the Language in Interaction Consortium from Netherlands Organization for Scientific Research (NWO). OJ was supported by James S. McDonnell Foundation Understanding Human Cognition Collaborative Award [220020448] and the Royal Society Wolfson Research Merit Award. LD was supported by the European Research Council (grant #773079-CoAct awarded to J.Holler). ES was supported by NWO (Veni grant 016.Veni.198.065). We are very grateful to Nick Wood, for helping us in editing the video stimuli, and to Gina Ginos, for being the actress in the videos.

